# Multiple mating rescues offspring sex ratio but not productivity in a haplodiploid exposed to developmental heat stress

**DOI:** 10.1101/2022.04.07.487407

**Authors:** Sofia G Costa, Sara Magalhães, Leonor R Rodrigues

## Abstract

1. Reproduction is generally more sensitive to high temperatures than survival and arguably a better predictor of the response of populations to climate change than survival estimates. Still, how temperature simultaneously impacts male and female reproductive success, the mating system and the operational sex ratio remains an open question.
2. Here, we addressed how a sublethal high temperature affects the reproductive system of the haplodiploid spider mite *Tetranychus urticae*. Males and females maintained at 25°C or 36°C during development were paired and the fertility of both sexes, their mating and remating eagerness, and the paternity of the offspring of females with different mating histories were measured.
3. Female and male fertility decreased at 36°C compared to 25°C, resulting in lower offspring production and a more male-biased sex ratio, respectively, because of haplodiploidy. However, when females remated, the pattern of first male sperm precedence typically seen in this species was disrupted, with more than one male contributing to the offspring. This was accompanied by reduced mating eagerness in pairs with partially sterile males and increased remating eagerness in pairs in which at least one sex was partially sterile in the first mating.
4. The observed temperature-induced changes in pre- and post-copulatory traits allowed restoring the offspring sex ratio but did not lead to the recovery of offspring number.
5. Our results show that temperature induces changes in the spider mite mating behaviour and mating system, with potential impact on sexual selection and sexual conflict. However, such changes may not be sufficient to buffer the impact of extreme temperatures on their populations.

## Introduction

Climate change is causing average local temperatures to rise and a higher frequency of heatwaves (Buckley & Huey, 2016; Kingsolver et al., 2013). These changes in the global thermal environment affect the biology of organisms at several levels, from behavioral and life-history traits of individuals to geographic range and abundance of populations, thereby posing a severe threat to biodiversity (Kellermann et al., 2012; Parmesan, 2006). This may be particularly detrimental to ectotherms, which represent the vast majority of terrestrial biodiversity (Stork et al., 2015), as their basic physiology is dependent on ambient temperatures (Hochachka & Somero, 2002).

Although viability has been often used as an indicator of species thermal limits (Geerts et al., 2015; Kellermann et al., 2012), it is becoming increasingly clear that thermal sensitivity differs among the main life-history traits, with male and female fertility being often the most sensitive traits in many taxa (Ma et al., 2020; Ma et al., 2015; Zhang et al., 2015; Zhao et al., 2014). Therefore, fertility is expected to be a better predictor of the response of populations to climate change than survival estimates, a prediction that is corroborated by recent studies (Parratt et al., 2021; van Heerwaarden & Sgrò, 2021; Walsh et al., 2019). Addressing the effect of these sublethal temperatures on reproductive traits is thus key to understand the impact of climate change on biodiversity.

Often, responses to sublethal high temperatures are sex-specific (Iossa, 2019), with important evolutionary consequences. For instance, in several species, males are more affected by sterility than females, with male-induced sterility leading to females becoming more eager to mate to compensate for less sperm being transferred. Most of these studies on thermally-induced sterility concern polyandrous species (Baur et al., 2022; Berger et al., 2011; Vasudeva et al., 2019), with a single study using a monandrous species (Sutter et al., 2019). This is unfortunate because in monandrous species, or species with single paternity, thermally-induced multiple paternity implies a change in the mating system, that in turn should result in a rearrangement of the selection pressures felt by males and females (García-Roa et al., 2020). Differential responses to sublethal high temperature between the sexes can also result in cryptic shifts in the operational sex ratio (OSR), as observed in some insect species (García-Roa et al., 2020), including in *Drosophila virilis*. In this species, male, but not female, sexual maturation is delayed at high temperature, with the operational sex ratio of populations becoming more female-biased without observable changes on the adult sex ratio (Walsh et al., 2021). Alternatively, changes in OSR can be accompanied by changes in adult sex ratio (ASR) in species with thermal-dependent sex determination and in haplodiploids (Maurer et al., 2021; Mobley & Jones, 2009). Specifically, in haplodiploids, female sterility results in reduced offspring production but male sterility leads to a male-biased ASR, because fertilized eggs only generate (diploid) daughters, while (haploid) sons are generated from unfertilized eggs (Helle & Sabelis, 1985). In these cases, heat-induced sterility could result in changes in both sex ratios and productivity. Yet, no study to date investigated the effect of sublethal temperatures on such species, despite the recognized impact of these variables on population dynamics and their role as drivers of plastic and/or evolutionary changes (Geffroy & Wedekind, 2020; Schmickl & Karsai, 2010).

Here, we study the effect of high sublethal temperatures on male and female fertility, paternity share and mating behavior in the spider mite *Tetranychus urticae*. Spider mites have a haplodiploid sex-determination system (Helle & Sabelis, 1985), typically with a female-biased ASR in the laboratory (ca. 70-80% females; Mitchell, 1972). This species presents a pattern of nearly complete first-male sperm precedence (Helle, 1967; Rodrigues et al., 2020) with mated females staying out of the mating pool, hence their OSR is often male-biased. The costs of male sterility are thus likely to be high. The mating behavior of spider mites follows the expected behavior of species with this sperm precedence pattern: males actively guard immature females to mate with them as soon as they reach adulthood (Potter et al., 1976), they prefer virgin over mated females (Oku, 2014; Rodrigues et al., 2017) and have prolonged copulations to ensure sperm precedence (Satoh et al., 2001). Although the response of spider mites to temperature has been vastly documented (Farazmand, 2020; Gotoh et al., 2010; Praslička & Huszár, 2004; Riahi et al., 2013; Zou et al., 2018), to date no study has tested if the sperm precedence pattern in spider mites can be disrupted, and the behavior altered, in response to heat-induced sterility.

## Materials and Methods

### Spider mite populations and rearing conditions

Two populations of spider mites (red form of *Tetranychus urticae*), one resistant and one susceptible to the pesticide etoxazole, were used in the experiments. These two populations shared the same genetic background, except for the presence or absence of a single nucleotide recessive mutation on the chitin synthase 1 gene (CHS1) causing resistance to etoxazole (van Leeuwen et al., 2012). Briefly, the procedure used to create these two populations was as follows. Both populations originated from the same five population replicates (the ‘iC’ populations in Rodrigues et al., 2022), themselves formed by merging six *Wolbachia-infected*, homozygous etoxazole-susceptible, populations collected in the region of Lisbon, Portugal, in 2013 (Zélé et al., 2018). Five etoxazole-resistant population replicates were then created by performing eight generations of backcross between each of the susceptible population replicates, and a homozygous etoxazole-resistant paternal line *SB9.Rif* (originally collected in Crete, Greece, in 2006; van Leeuwen et al., 2012). Each generation of backcross was followed by a generation of selection (with etoxazole exposure) of the haploid males. This procedure allowed the complete introgression of the etoxazole resistant allele into the susceptible population replicates, but with complete retention of their mitochondrial genome and an estimated retention of 99.6% of their nuclear genome. The presence or absence of mutation of the CHS1 gene in all susceptible and resistant population replicates was confirmed by PCR-RFLP (as described in van Leeuwen et al., 2012). All population replicates were then treated with rifampicin following the procedure described in Zélé et al. (2020). Both susceptible and resistant replicates were merged in September 2020 to create, respectively, the susceptible (*Wu.SS*, uninfected with Wolbachia, susceptible to the pesticide) and resistant (*Wu.RR*, uninfected with Wolbachia and resistant to the pesticide) base populations used here.

The two base populations were maintained in large numbers (> 2000) under continuous generations on entire plants from September 2020 to January 2021, then in smaller numbers (> 1000) and with discrete generations from January 2021 onwards (to facilitate obtaining individuals of a specific stage for the experiments). These populations were kept on bean leaves (*Phaseolus vulgaris*, variety Contender, provided by Germisem, Portugal) under controlled conditions (25°C; 16h Light: 8h Dark photoperiod). Bean plants were germinated and grown in an isolated and herbivore-free room for 14 days under controlled conditions (16h Light (at 25°C): 8h Dark (at 20°C) photoperiod) before being exposed to spider mites.

### Experimental setup

#### Fertility and paternity share

To test if the offspring paternity share is altered when the first male has reduced fertility due to developmental heat stress, we crossed a resistant female (from *Wu.RR*) with a susceptible male (from *Wu.SS*) and then with resistant males (from *Wu.RR*), following a protocol developed in Rodrigues et al., (2020). Paternity share was determined by exposing the resulting offspring to the pesticide etoxazole. This pesticide inhibits the synthesis of chitin, preventing eggs and juveniles from hatching and molting, respectively, and resistance is determined by a single, recessive, allele (van Leeuwen et al., 2012). Because spider mites are haplodiploid and females always came from the resistant population, the pesticide application will only affect daughters (diploids), while sons (haploids), which only inherit the genetic material of their mothers, will always be resistant. Thus, when exposed to pesticide during development, the offspring will consist of alive sons, alive daughters (sired by resistant males) and/or unhatched eggs (sired by susceptible males).

Four treatments, differing in the temperatures under which females and their first mate developed, were tested: pairs where both individuals developed at control temperature (25°C; ♀25 x ♂25), pairs where both individuals were exposed to heat stress during development (36°C; ♀36 x ♂36), and pairs where only the male or the female was exposed to heat stress (♀25 x ♂36 and ♀36 x ♂25, respectively). These treatments were orthogonally crossed with mating history treatments, in which females from all temperature treatments mated once, twice or multiple times. The second and subsequent matings always involved control males.

To obtain females from all temperature treatments and mating histories, virgin males and females were taken from cohorts of each base population kept at either 25 or 36°C (control temperature and heat stress, respectively). Five resistant virgin females and 5 susceptible virgin males were placed on a mating disc (bean discs of 0.5 cm^2^ placed inside a Petri dish with water-saturated cotton) at 25°C to mate for 30 minutes. Once mated, males were discarded, and females, in groups of 5, were either placed on leaf discs of 0.95 cm^2^ (Single Mated females, SM) or transferred to a new mating disc (0.5 cm^2^) with 5 resistant virgin males developed at control temperature and given 2 hours to remate (Double Mated females, DM). Again, once remated, females were placed in groups of 5 on leaf discs of 0.95 cm^2^, while males were discarded after their first mating. Both matings were observed in order to verify that copulation occurred, but no data was recorded. A subset of the SM females was transferred in groups of 5 to a new disc (0.95 cm^2^) with 5 resistant virgin males that developed at control temperature, where they could freely remate multiple times and/or with multiple males for 24 hours (Multiply Mated females, MM). The next day, single-, double- and multiply-mated females were individually transferred to new discs (2.55 cm^2^) placed on etoxazole-soaked cotton (diluted in water at 0.5g/L). Females had 4 days to oviposit, after which they were discarded, and the eggs were counted. The survival status of each female was recorded every day until female removal. The number of hatched and unhatched eggs was recorded 8 days after oviposition started. Adult offspring was sexed, counted and removed 12 or 14 days after the onset of oviposition.

This experiment was divided in 4 blocks, 2 per week in consecutive days, each including all treatments. SM females that did not mate and DM females that did not remate were discarded. Replicates with damaged females (i.e., females that were hurt during manipulation) were excluded from the analysis. For each female mating history (SM, DM and MM), 27 to 43 replicates were analyzed per temperature treatment.

#### Mating behavior

To determine if female mating and remating eagerness were affected by developmental heat stress, four mating treatments, targeting focal females and their first mate, were established: pairs where both individuals developed at control temperature (25°C; ♀25 x ♂25), pairs where both individuals were exposed to heat stress during development (36°C; ♀36 x ♂36), and pairs where only the male or the female was exposed to heat stress (♀25 x ♂36 and ♀36 x ♂25, respectively). Individuals were collected from cohorts of the base population *Wu.RR* kept at 25 or 36°C (control temperature and heat stress, respectively).

Virgin females and males were paired on a bean disc (0.5 cm^2^) placed in a Petri dish with water-saturated cotton and given 30 minutes to copulate. The latency to copulation and its duration were recorded using the app Stopwatch: StopNow Free. Once mated, females were either individually placed on leaf discs of 0.95 cm^2^ (Single Mated females; SM) or transferred to a new disc (0.5 cm^2^) to which five virgin males that had developed at 25°C were added. Females were then given 2 hours to remate, with males being replaced after each copulation. The latency to each copulation and its duration were recorded. Afterwards, these females were individually placed on leaf discs of 0.95 cm^2^ (Multiple Mated females; MM). The next day, both SM and MM females were individually transferred to larger leaf discs (2.55 cm^2^), and their offspring was monitored as above.

This experiment was performed in 8 blocks, 2 per week in consecutive days, each including all treatments. Individuals were maintained at 25°C during the mating observations and afterwards. Replicates with single mated females that did not mate, multiply mated females that did not remate or damaged females (i.e., females that were hurt during manipulation) were excluded from the analysis. 89 to 93, and 15 to 43 replicates, were analyzed per treatment to determine the effect of developmental heat stress on female mating and remating eagerness, respectively.

### Statistical analyses

All statistical analyses were performed using the software R (version 4.0.3; R Core Team, 2020). We performed generalized mixed-effects models (GLMM) and linear mixed-effects models (LMM) implemented in *lme4* (Bates et al., 2015) and *glmmTMB* (Brooks et al., 2017), depending on the data and error structure. Maximal models were simplified by removing non-significant terms (*p* < 0.05) from the highest- to the lowest-order interaction (Crawley, 2012; Tables S1 and S2). The significance of each explanatory variable was determined by chi-squared tests for discrete distributions, and Wald F tests for continuous distributions (Bolker et al., 2009). *A posteriori* contrasts with Bonferroni corrections were made to interpret the effect of factors with more than two levels, using the *emmeans* package (Length, 2020; Tables S3 and S4). Graphic representations of the data were produced with the software package *ggplot2* (Wickham, 2016).

#### Fertility and paternity share

To test how developmental heat stress affected fertility and paternity share, three variables were analyzed (Table S1): total offspring number (i.e., the number of daughters, sons and unhatched eggs per female), as a measure of female fertility, the contribution of all males to the offspring (i.e., the number of daughters and unhatched eggs, over the total offspring), and the contribution of the second and subsequent males to the offspring (i.e., the proportion of daughters over the fertilized offspring).

The total offspring number (unhatched eggs + daughters + sons) was analyzed using a GLMM with a quasi-Poisson error distribution (package *glmmTMB*). The contribution of all males to the offspring was computed using *cbind* (with the sum of daughters and unhatched eggs and the number of sons as arguments), a beta-binomial error distribution and a parameter to account for zero inflation (ziformula ~1; package *glmmTMB*). The contribution of the second and subsequent males to the offspring was computed using the function *cbind* with the number of daughters and the number of unhatched eggs as arguments and analyzed using a GLMM with a binomial error distribution (package *lme4)*.

The developmental temperature of the female (25°C or 36°C) and of the first male (25°C or 36°C), the female mating history (SM, DM or MM), as well as their interaction, were added as fixed factors in the models for the total number of offspring and for the contribution of all males to the offspring. In the model for the contribution of the second and subsequent males to the offspring, the developmental temperature of the pair (female and first male; ♀25 x ♂25, ♀25 x ♂36, ♀36 x ♂25 and ♀36 x ♂36), the female mating history (SM, DM or MM) and their interaction were included as fixed factors. The developmental temperature of the pair was used instead of the developmental temperatures of the female and of the first male separately to avoid convergence issues. As expected, when females were single mated, the proportion of daughters (i.e., the contribution of the second and subsequent males) was zero, with only one replicate having a proportion of daughters above zero (in treatment ♀25 x ♂36; proportion = 0.034). To deal with this lack of variance that impeded the statistical analysis, we manually added one replicate to the SM mating history of each treatment: the values of these replicates were equal to the one replicate above zero in treatment ♀25 x ♂36, this way solving the variance problem while maintaining the original differences among treatments. Block, temperature and humidity during mating observations were included as random factors in all models (Table S1).

#### Mating behavior

To analyze how mating and female remating eagerness were affected by developmental heat stress, we analysed latency to the first copulation and the duration of that copulation, the latency to the second copulation and the duration of that copulation (Table S2). The number of adult offspring and sex ratio were also analyzed to infer female and male fertility, respectively, and to confirm replicability with the results obtained in the previous experiment (cf. Supplementary Information 1).

The latency to the first and second copulations and their duration were analyzed using LMMs with a gaussian error distribution (package *lme4*). The data was boxcox transformed (latency and duration of first copulation: λ= 0.39; latency of second copulation: λ= 0.11; duration of second copulation: λ=-0.05; *MASS* package; Venables & Ripley 2002) to improve the model fit.

In all models, block and temperature and humidity during mating observations were included as random factors, and the developmental temperature of the female (25°C or 36°C) and of the first male (25°C or 36°C), as well as their interaction, were added as fixed factors. Additionally, the mating rate was included as a covariate in the models for the latency to the second copulation and its duration, as these traits might be affected by female eagerness to remate (Table S2).

## Results

### Fertility and paternity share

The total number of offspring was affected by the interaction between the developmental temperature of females and of their first mate (χ^2^1 = 4.608, *p* = 0.032; Table 1), but not by the female mating history (χ^2^2 =1.782, *p* = 0.410; Table 1). Indeed, thermally stressed females produced ca. 74% fewer offspring than control females (9±0.52 at 36°C and 35±0.84 at 25°C), independently of the developmental temperature of their mates or their mating history (Table S3; Figure 1a). In addition, control females produced ca. 15% fewer offspring when first mating with thermally stressed males (32±0.98) than when mating with control males (38±1.32; Table S3; Figure 1a).

**Figure 1.**
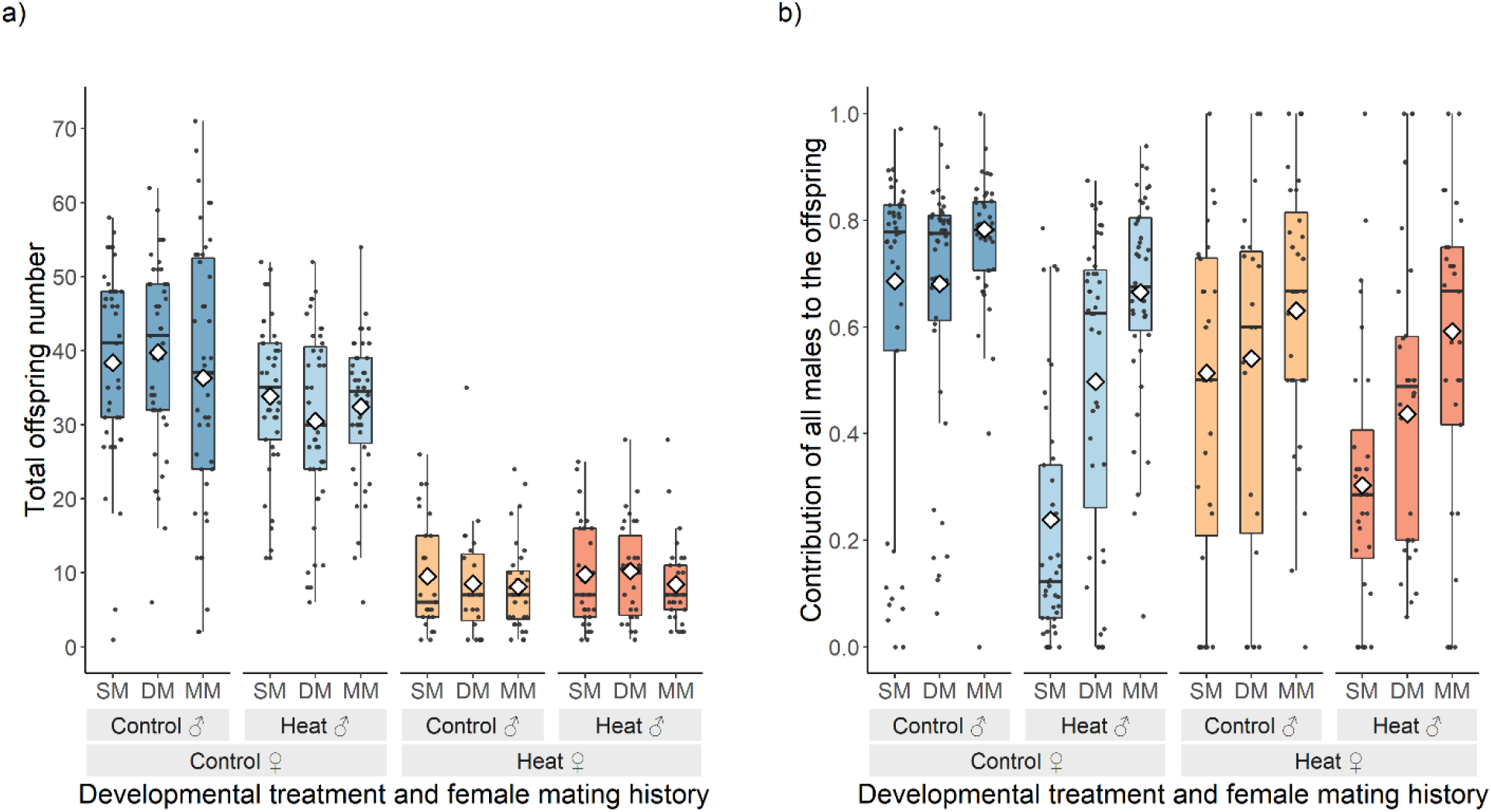
Effect of high developmental temperature on fertility and the contribution of males to the offspring. Females mated once (SM), twice (DM) or multiply over 24 hours (MM). The first mating involved a male susceptible to the pesticide etoxazole and developed at the temperature described in the figure, while all subsequent matings involved control males resistant to etoxazole. a) Total number of offspring, including sons, daughters and unhatched eggs, representing the fertility of the female; b) Contribution of all males to the offspring, represented by the proportion of fertilized offspring. Boxplots display a median line, interquartile range (IQR) boxes, 1.5*IQR whiskers and data points. Colors dark blue, light blue, light orange and dark orange represent treatments where both individuals were developed at the control temperature, only the male or the female were heat-stressed, and both individuals were heat-stressed, respectively. White diamonds represent the mean values per conditions tested.

**Table 1.** Statistical results of the effect of temperature on fertility and paternity share. “Df”: degrees of freedom. “χ^2^”: Chi-square value obtained in each analysis. “Total offspring number”: number of daughters, sons and unhatched eggs per female; “Contribution of all males to the offspring”: number of daughters and unhatched eggs, over the total offspring; “Contribution of the second and subsequent males to the offspring”: proportion of daughters over the fertilized offspring. “♀ temperature”: developmental temperature of the females tested; “♂ temperature”: developmental temperature of the first male that mated with a female; “♀ mating history”: the number of times a female mated; “Temperature of pair”: developmental temperatures of the female and the first male crossed in each replicate. Statistically significant terms (*p* ≤ 0.05) are represented in bold.

| Dependent var. | Explanatory var. | Df | $\chi^2$ | P-value |
| --- | --- | --- | --- | --- |
| Total offspring number | ♀ **temperature** | **1** | **269.271** | **<0.001** |
|  | ♂ **temperature** | **1** | **7.376** | **0.006** |
|  | ♀ mating history | 2 | 1.782 | 0.410 |
|  | ♀ **temperature** * ♂ **temperature** | **1** | **4.608** | **0.032** |
|  | ♀ temperature * ♀ mating history | 2 | 0.032 | 0.984 |
|  | ♂ temperature * ♀ mating history | 2 | 2.877 | 0.237 |
|  | ♀ temperature * ♂ temperature * ♀ mating history | 2 | 2.477 | 0.290 |
| Contribution of all males to the offspring | ♀ **temperature** | **1** | **12.420** | **<0.001** |
|  | ♂ **temperature** | **1** | **11.543** | **<0.001** |
|  | ♀ **mating history** | **2** | **7.248** | **0.027** |
|  | ♀ **temperature** * ♂ **temperature** | **1** | **4.510** | **0.034** |
|  | ♀ temperature * ♀ mating history | 2 | 3.138 | 0.208 |
|  | ♂ **temperature** * ♀ **mating history** | **2** | **27.528** | **<0.001** |
|  | ♀ temperature * ♂ temperature * ♀ mating history | 2 | 5.343 | 0.069 |
| Contribution of the second and subsequent males to the offspring | **Temperature of pair** | **3** | **516.785** | **<0.001** |
|  | ♀ **mating history** | **2** | **53.679** | **<0.001** |
|  | **Temperature of pair** * ♀ **mating history** | **6** | **22.956** | **<0.001** |

The contribution of all males to the offspring was significantly affected by the interaction between the developmental temperature of the first male and the mating history of the female (χ^2^2 = 27.528, *p* =<0.001; Table 1) and between the developmental temperature of the female and of her first mate (χ^2^1 = 4.510, *p* = 0.034; Table 1). As expected, in control males, there was no significant effect of mating history, with the proportion of fertilized offspring averaging 0.70±0.03 (Table S3; Figure 1b). Moreover, the proportion of fertilized offspring decreased when the female and/or the first male she mated with were exposed to heat stress. Indeed, the proportion of fertilized offspring of females that mated once with heat-stressed males was reduced by 0.41±0.08 compared to that of females mated once with control males (Table S3; Figure 1b). However, when these females were allowed to remate, the proportion of fertilized offspring increased, being the highest when females mated multiply (Table S3; Figure 1b), in which case it reached values close to those of females mated with control males.

The contribution of the second and subsequent males to the offspring was significantly affected by the interaction between the female mating history and the developmental temperature of the pair (χ^2^6 = 22.956, *p* =<0.001; Table 1). As expected, the proportion of offspring sired by the second or subsequent males was the lowest when the females and their first mates developed at control temperature (Table S3; Figure 2). When at least one of the individuals of the first mating was exposed to developmental heat stress, the paternity share of subsequent males increased significantly. Indeed, when the first male to mate with a female was exposed to heat stress, second and subsequent males contributed the most to the offspring, especially if the female developed at control temperature (Table S3; Figure 2). In addition, a significant increase in the proportion of daughters when females mated multiply, compared to when they mated twice, was observed only in control females (Table S3; Figure 2).

**Figure 2.**
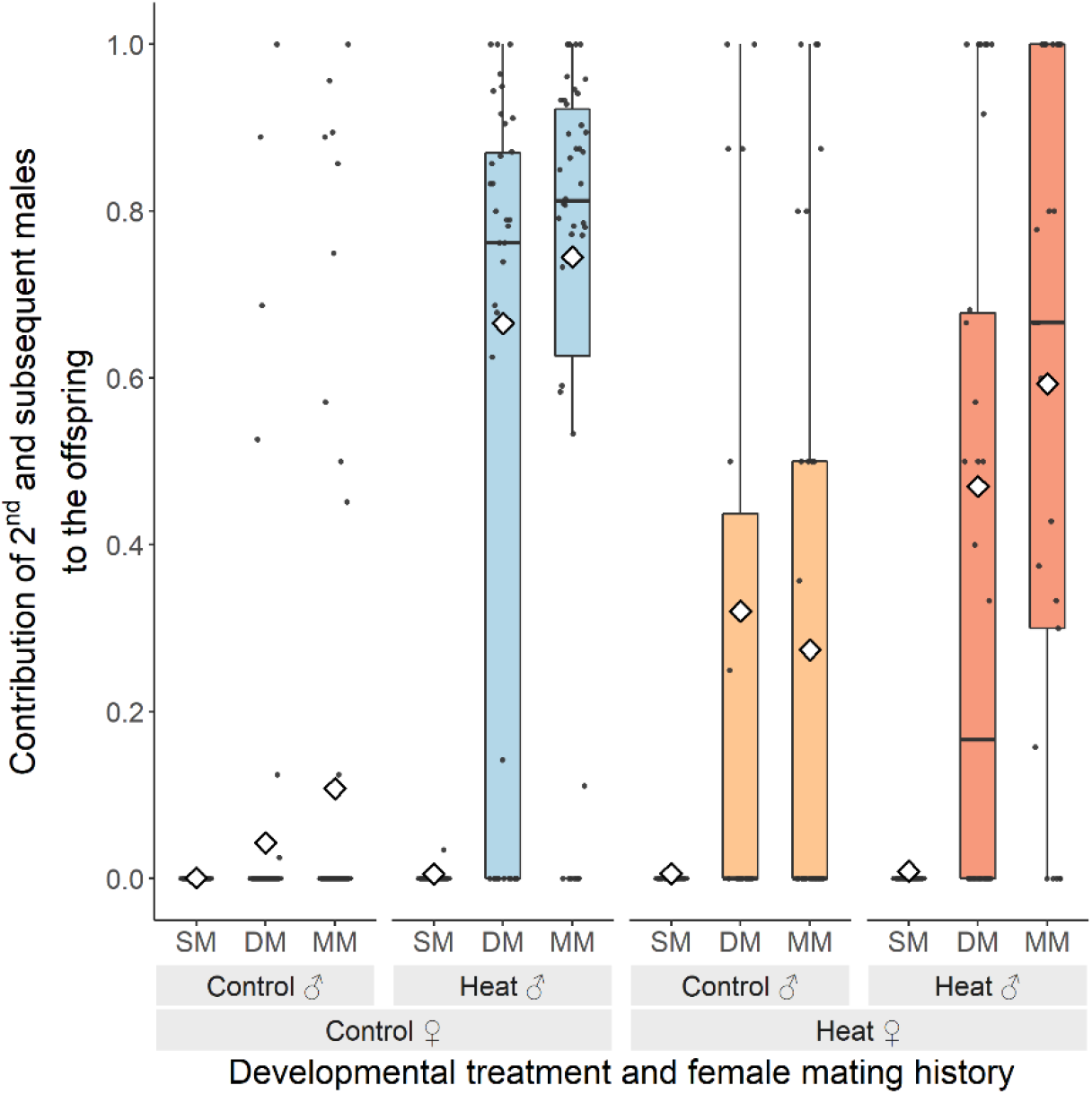
Effect of high developmental temperature on sperm priority. Females developed at 25°C or 36°C mated once (SM), twice (DM) or multiply over 24 hours (MM). The first mating involved a male susceptible to the pesticide etoxazole and that developed at the temperature described in the figure, while all subsequent matings involved control males resistant to etoxazole. The contribution of second matings (and subsequent matings, in the case of MM) to the offspring is represented by the proportion of daughters (resist to pesticide). Boxplots display a median line, interquartile range (IQR) boxes, 1.5*IQR whiskers and data points. Colors dark blue, light blue, light orange and dark orange represent treatments where both individuals were developed at the control temperature, only the male or the female were heat-stressed, and both individuals were heat-stressed, respectively. White diamonds represent the mean values per conditions tested.

### Mating behavior

The interaction between the developmental temperature of both sexes affected the latency to the first copulation (F_1,342.51_ = 4.647, *p* = 0.032; Table 2). Indeed, heat-stressed males took on average the double of the time to start copulating with control females as compared to control males and ca. 30% longer to mate with heat-stressed females as compared to control males (Figure 3a). The duration of the first copulation was only affected by the developmental temperature of the male (F_1,344.62_ = 52.403, *p* =<0.001; Table 2), with pairs including heat-stressed males having ca. 40% shorter copulations than pairs including control males (Figure 3b).

**Figure 3.**
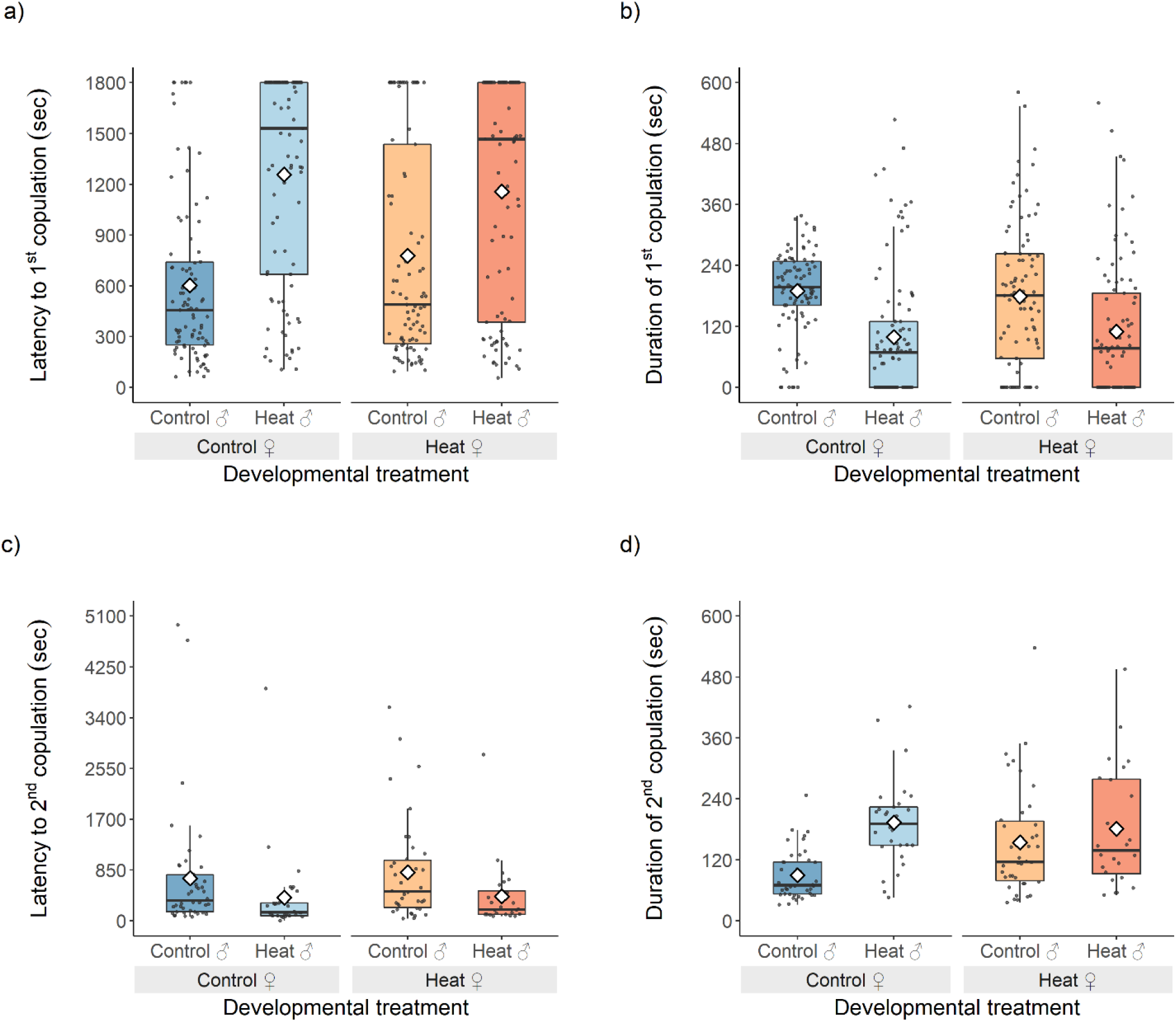
Effect of high developmental temperature on mating behavior. Females mated once (SM) or multiply over 2 hours (MM). The first mating involved a male developed at the temperature described in the figure, while the subsequent matings involved control males. Latency to mate (a), duration of first copulation (b), latency to remate (female’s second mating (c)) and duration of second copulation (d) are displayed in seconds. Boxplots display a median line, interquartile range (IQR) boxes, 1.5*IQR whiskers and data points. Colors dark blue, light blue, light orange and dark orange represent treatments where both individuals were developed at the control temperature, only the male or the female were heat-stressed, and both individuals were heat-stressed, respectively. White diamonds represent the mean values per conditions tested.

**Table 2.**
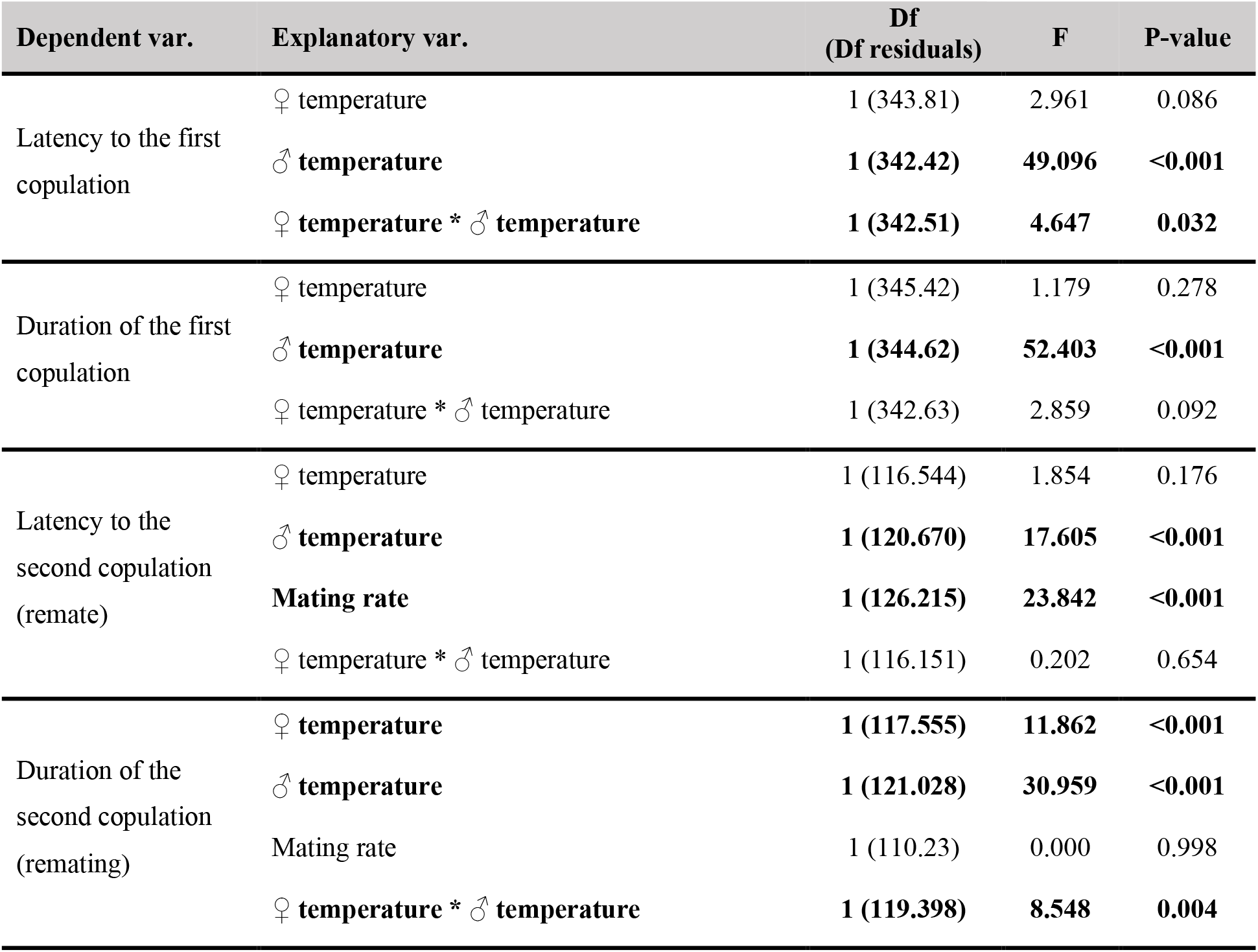
Statistical results of the effect of temperature on mating behavior. “Df”: degrees of freedom. “F”: sum of squares obtained from the F-test. “Latency to the first copulation”: time elapsed from pairing until the first mating of a female in seconds; “Duration of the first copulation”: time elapsed from the beginning until the end of the first mating of a female in seconds; “Latency to the second copulation (remate)”: time elapsed from pairing until second mating of a female in seconds; “Duration of the second copulation (remating)”: time elapsed from the beginning until the end of the second mating of a female in seconds. “♀ temperature”: developmental temperature of the females tested; “♂ temperature”: developmental temperature of the first male that mated with a female; “Mating rate”: number of matings done by multiple mated females; “♀ mating history”: the number of times a female mated. Statistically significant terms (*p* ≤ 0.05) are represented in bold.

The latency to the second copulation with a fertile male was affected by the developmental temperature of the first male to mate with the female (F_1,120.670_ = 17.605, *p* =<0.001; Table 2), with pairs including females first mated with heat-stressed males taking more than half of the time to start remating than pairs with females first mated with control males ( Figure 3c). The duration of this second copulation was affected by the interaction between the developmental temperature of the female and of her first mate (F_1,119.398_ = 30.959, *p* =<0.001; Table 2): control females first paired with control males had the shortest copulation duration, with all other pairs involving at least one individual exposed to heat stress showing a larger duration (Table S4; Figure 3d).

## Discussion

In this study we showed that at high sublethal developmental temperature, not only was the fertility of both sexes hindered in spider mites, but the paternity share was altered, with males other than the first siring offspring. In accordance with the toll on fertility and change in paternity share, matings involving males that developed at high temperature took longer to occur and were shorter than matings with fertile males, while females involved in matings with partially sterile males were also more eager to remate with fertile males and rematings involving partially sterile individuals lasted longer. These changes resulted in a nearly total restoration of the proportion of fertilized offspring (i.e., daughters) but not in recovery of the number of offspring produced.

A reduction in fertility with high temperature has been previously demonstrated in many taxa for both sexes (Walsh et al., 2019), with accumulating evidences of deficiencies, unviability, depletion or incomplete transfer of sperm/ejaculate in males (Gillooly et al., 2001, 2002; Hurley et al., 2018; Lucio et al., 2016; Nguyen et al., 2013; Rahman et al., 2018; Rao et al., 2016; Sales et al., 2018; Setchell, 1998; Zeh et al., 2014) and changes in egg size and oocyte degeneration in females (Hansen, 2009; Paxton et al., 2016). Curiously, although spider mites are important crop pests known to be affected by temperature (Bayu et al., 2017; Kasap, 2004; Praslička & Huszár, 2004; Riahi et al., 2013), no study so far had disentangled the effects of heat stress on males and females in this group. By filling this gap, we found that male sterility resulted mostly in a shift in adult sex ratio towards more males and a slight change in offspring number. In line with this, matings with thermally stressed males showed increased latency to copulation and decreased mating duration, compared to matings with control males, suggesting that females avoid mating with males exposed to developmental high temperature and/or that such exposure affects male mobility or their perception of the presence of a female. A previous study in *T. urticae* has found similar behavioural changes in females involved in heterospecific matings that resulted in no fertilized offspring (Clemente et al., 2016), suggesting similar behaviors occur with different sources of infertility. In turn, heat stress in females translated simultaneously into reduced offspring number and a slight change in adult sex ratio towards more males. In this case, no change in mating eagerness was observed between matings involving control *vs* heat stressed females, again in line with the results of Clemente et al. (2016) that found a lack of male preference for conspecific *vs* heterospecific females. Interestingly, when both sexes were exposed to heat stress, the adult sex ratio of the offspring was equivalent to that observed when only the male was heat stressed, and the number of adult offspring was equivalent to that found when only the female was exposed to heat, suggesting the effect of temperature in both sexes is the sum of the independent effect of temperature in males and females, respectively.

The effect of sublethal high temperature in males could have a snowball effect, as male partial sterility results in the production of more (partially sterile) males. Moreover, the pattern of sperm precedence of this species is such that the first male generally sires all the offspring of a female, suggesting that females engaging in matings with partially sterile males might pay the high cost of not producing fertilized eggs. The combination of these two phenomena is expected to rapidly lead to a highly male-biased sex ratio in the population. Adding to this the fact that females exposed to heat stress produced fewer offspring, the probability of population collapse due to this sublethal temperature seems to be high. This is particularly relevant because spider mites occur in fragmented populations in which females are the dispersing sex, hence the number of females produced can be used as a measure of fitness (Metz & Gyllenberg, 2001).

The reduction in male and female fertility due to developmental heat stress has, however, led to the fertilization of some eggs by males other than the first, resulting in an overall higher proportion of fertilized viable offspring when females mated more than once. This recovery of the fertilized offspring of females almost reached the values of crosses only involving individuals exposed to optimal temperatures, being more pronounced when females mated with more males, as found in spiders (Matzke et al., 2022). In addition, females that mated first with thermally stressed males mated faster and showed an increase in mating duration when remating with control males, compared to females that had first mated with control males, suggesting females modulate their mating investment. All in all, the pre- and post-copulatory behavior observed could help preventing population collapse, provided that females encounter fertile males via dispersal to, or migration from, cooler places. Two other studies in *Drosophila pseudoobscura* and *Tribolium castaneum* (Sutter et al., 2019; Vasudeva et al., 2021) have found similar changes in mating behavior coupled with benefits of multiple mating after exposure to sublethal high temperature, whereas several others have found benefits of multiple mating against other sources of male sterility (Friesen et al., 2014; Gavriel et al., 2009; Kraaijeveld & Chapman, 2004; Landeta-Escamilla et al., 2016). This suggests that females can make dynamic remating decisions with multiple mating functioning as a mechanism against different sources of mating failure or male infertility. This can help explain why multiple mating is so prevalent across species, including in those in which first male sperm precedence is the norm, like spider mites.

Unlike what was observed for the sex ratio and what has been documented in diploids (Sutter et al., 2019; Vasudeva et al., 2021), the fecundity of heat-stressed spider mites did not recover with multiple mating. This suggests that in haplodiploids, multiple mating improves egg fertilization but not egg production. Furthermore, in heat-stressed females, the recovery of offspring sex ratio was lower, suggesting that these females are unable to fully store or use the sperm they receive. High temperature has been shown to affect egg size in females (Vasudeva et al., 2019) which in spider mites mediates sex allocation (Macke et al., 2012), giving a potential explanation for the inability to fully recover the offspring adult sex ratio. This might also explain why rematings with heat-stressed females lasted longer than those with control females. Whatever the mechanism at play, it seems that polyandry can be less beneficial to endure warming in haplodiploid species than in diploid species due to the decoupling of egg fertilization and egg production, begging the question of whether these species are particularly vulnerable to climate change.

Our findings have important consequences for our knowledge of the reproductive behaviour and the mating system of spider mites. Indeed, at optimal temperatures, the ASR of spider mite populations is typically female biased, with an average female proportion of circa 0.7 (Helle & Sabelis, 1985). In addition, due to nearly complete first male precedence, most females leave the mating pool as soon as they mature unlike males that remain in the mating pool, not becoming sperm depleted unless they mate with more than ca. 15 females a day (Krainacker & Carey, 1989). Hence, males and females have different potential reproductive rates (Parker & Simmons, 1996), leading to an OSR that is male-biased (Kokko et al., 2012). The bias in the OSR is expected to result in strong male-male competition (García-Roa et al., 2019) but this may not translate into strong sexual conflict, as neither males nor females benefit from matings involving mated females (Rodrigues et al., 2020). Under heat stress, in contrast, females will remain in the mating pool after the first mating as shown here, hence the OSR will approach the ASR. Female multiple mating will also lead to a steeper Bateman gradient in females, as they will benefit from matings after the first (Kvarnemo & Simmons, 2013). In turn, males will no longer fertilize all the offspring of females they mate with, so their gradient will become shallower. Together, these temperature-driven changes are expected to strongly modulate the expression of sexual selection and conflict in this species.

## Notes

### Competing Interest Statement

The authors have declared no competing interest.

