## Supplementary Material for "Multiple mating rescues offspring sex ratio but not productivity in a haplodiploid exposed to developmental heat stress"

### Table of contents

**Table S1 –** Description of the statistical models used for data analysis of the effect of temperature on fertility and paternity share ……………….…………………………………………………………….... 2

**Table S2 –** Description of the statistical models used for data analysis for the effect of temperature on mating behavior ……………………………………………………………………………………….. 3

**Table S3 –** *A posteriori* contrasts of significant explanatory variables for the effect of temperature on fertility and paternity share ………………………………………………………………………………. 4

**Table S4 –** *A posteriori* contrasts of significant explanatory variables for the effect of temperature on mating behavior ……………………………………………………………………………………….. 6

**Supplementary Information 1 –** The effect of temperature on the offspring number and sex ratio of females from the mating behavior experiment ..………………………………………………………….. 7

**Table S1 – Description of the statistical models used for data analysis of the effect of temperature on fertility and paternity share.** "Sample size": total number of replicates included in each analysis. "Maximal model": complete set of explanatory variables included in the model. "Minimal model": model containing only the variables that were statistically significant (round brackets indicate that the variable was included as a random factor). Square brackets indicate the error structure used ("b": binomial; "bbI": beta-binomial, accounting for zero inflation; "qp": quasi-Poisson). "Total offspring number": number of daughters, sons and unhatched eggs per female; "Contribution of all males to the offspring": number of daughters and unhatched eggs, over the total offspring; "Contribution of the second and subsequent males to the offspring": proportion of daughters over the fertilized offspring. "♂": number of sons; "♀": number of daughters. "♀ temperature": developmental temperature of the females tested; "♂ temperature": developmental temperature of the first male that mated with a female; "♀ mating history": the number of times a female mated; "Temperature of pair": developmental temperatures of the female and the first male crossed in each replicate; "Block": the day at which the individual replicates were started; "Room temperature": the temperature (°C) registered in the observation room at the time of each behavioral observation; "Room humidity": the humidity (%) registered in the observation room at the time of each behavioral observation. [a] one individual replicate was added to the SM mating history of each cross. These analyses only include replicates in which fecundity was above zero. Replicates with damaged females (i.e., females that were hurt during manipulation) were excluded from all the analyses.

| Dependent variable | Response variable | Sample size | Maximal model | Minimal model | R subroutine [error struct.] |
|---|---|---|---|---|---|
| Total offspring number | Number of offspring | 422 | ♀ temperature * ♂ temperature * ♀ mating history + (1\|Block) + (1\|Room temperature) + (1\|Room humidity) | ♀ temperature * ♂ temperature + (1\|Block) | glmmTMB (qp) |
| Contribution of all males to the offspring | cbind (♀ + unhatched eggs, ♂) | 422 | ♀ temperature * ♂ temperature * ♀ mating history + (1\|Block) + (1\|Room temperature) + (1\|Room humidity) | ♂ temperature * (♀ temperature + ♀ mating history) + (1\|Block) | glmmTMB (bbI) |
| Contribution of the second and subsequent males to the offspring | cbind (♀, unhatched eggs) | 426 [a] | Temperature of pair * ♀ mating history + (1\|Block) + (1\|Room temperature) + (1\|Room humidity) | Temperature of pair * ♀ mating history + (1\|Block) | glmer [b] |

**Table S2 – Description of the statistical models used for data analysis of the effect of temperature on mating behavior.** "Sample size": total number of replicates included in each analysis. "Maximal model": complete set of explanatory variables included in the model. "Minimal model": model containing only the variables that were statistically significant (round brackets indicate that the variable was included as a random factor). Square brackets indicate the error structure used ("qp": quasi-Poisson; "bbI": beta-binomial, accounting for zero inflation; "g": gaussian). "Latency to the first copulation": time elapsed from pairing until the first mating of a female in seconds; "Duration of the first copulation": time elapsed from the beginning until the end of the first mating of a female in seconds; "Latency to the second copulation (remate)": time elapsed from pairing until second mating of a female in seconds; "Duration of the second copulation (remating)": time elapsed from the beginning until the end of the second mating of a female in seconds; "Number of adult offspring": number of offspring that reached adulthood per female; "Sex ratio": proportion of adult daughters over the total adult offspring. "♂": number of sons; "♀": number of daughters. "♀ temperature": developmental temperature of the females tested; "♂ temperature": developmental temperature of the first male that mated with a female; "Mating rate": number of matings done by multiple mated females; "♀ mating history": the number of times a female mated; "Block": the day at which the individual replicates were started; "Room temperature": the temperature (°C) registered in the observation room at the time of each behavioral observation; "Room humidity": the humidity (%) registered in the observation room at the time of each behavioral observation. [a] only includes replicates in which females mated multiply (mating rate > 1); [b] includes single and multiply mated females in which fecundity was above zero. No replicates with damaged females (i.e., females that were hurt during manipulation) were used in the analysis.

| Dependent variable | Response variable | Sample size | Maximal model | Minimal model | R subroutine [error struct.] |
|---|---|---|---|---|---|
| Latency to the first copulation | Latency to 1st mating | 361 | ♀ temperature * ♂ temperature + (1\|Block) + (1\|Room temperature) + (1\|Room humidity) | ♀ temperature * ♂ temperature + (1\|Block) | lmer [g] |
| Duration of the first copulation | Duration of 1st mating | 361 | ♀ temperature * ♂ temperature + (1\|Block) + (1\|Room temperature) + (1\|Room humidity) | ♂ temperature + (1\|Block) | lmer [g] |
| Latency to the second copulation (remate) | Latency to 2nd mating | 133 [a] | ♀ temperature * ♂ temperature + Mating rate + (1\|Block) + (1\|Room temperature) + (1\|Room humidity) | ♂ temperature + Mating rate + (1\|Block) | lmer [g] |
| Duration of the second copulation (remating) | Duration of 2nd mating | 133 [a] | ♀ temperature * ♂ temperature + Mating rate + (1\|Block) + (1\|Room temperature) + (1\|Room humidity) | ♀ temperature * ♂ temperature + (1\|Block) | lmer [g] |
| Number of adult offspring | Number of adult offspring | 204 [b] | ♀ temperature * ♂ temperature * ♀ mating history + (1\|Block) + (1\|Room temperature) + (1\|Room humidity) | ♀ temperature + (1\|Block) | glmmTMB (qp) |
| Sex ratio | cbind (♀, ♂) | 204 [b] | ♀ temperature * ♂ temperature * ♀ mating history + (1\|Block) + (1\|Room temperature) + (1\|Room humidity) | ♀ temperature + ♂ temperature * ♀ mating history + (1\|Block) | glmmTMB (bbI) |

**Table S3 – *A posteriori* contrasts of significant explanatory variables for the effect of temperature on fertility and paternity share.** *A posteriori* contrasts with Bonferroni corrections were done to interpret the significant effect of the fixed factors. "T ratio": the T-test value obtained in each comparison. "Total offspring number": number of daughters, sons and unhatched eggs per female; "Contribution of all males to the offspring": number of daughters and unhatched eggs, over the total offspring; "Contribution of the second and subsequent males to the offspring": proportion of daughters over the fertilized offspring. Comparisons were made between females developed at 25°C or 36°C ("♀25" and "♀36"), first mated with males developed at 25°C or 36°C ("♂25" and "♂36") and that mated once (SM), twice (DM) or multiply (MM). Statistically significant terms ($p \leq 0.05$) are represented in bold. *represents marginally significant values.

| Dependent var. | Comparison | T ratio | P-value |
|---|---|---|---|
| Total offspring number | **♀25 x ♂25 – ♀25 x ♂36** | **2.716** | **0.028** |
| | **♀25 x ♂25 – ♀36 x ♂25** | **16.409** | **<0.001** |
| | ♀36 x ♂25 – ♀36 x ♂36 | -1.122 | 1.000 |
| | **♀25 x ♂36 – ♀36 x ♂36** | **14.341** | **<0.001** |
| Contribution of all males to the offspring | **♀25 x ♂25 – ♀25 x ♂36** | **7.526** | **<0.001** |
| | **♀25 x ♂25 – ♀36 x ♂25** | **3.524** | **0.002** |
| | ♀36 x ♂25 – ♀36 x ♂36 | 2.501 | 0.051* |
| | ♀25 x ♂36 – ♀36 x ♂36 | 0.524 | 1.000 |
| | SM ♂25 – DM ♂25 | -0.415 | 1.000 |
| | SM ♂25 – MM ♂25 | -2.506 | 0.139 |
| | DM ♂25 – MM ♂25 | -2.079 | 0.420 |
| | **SM ♂25 – SM ♂36** | **7.625** | **<0.001** |
| | DM ♂25 – DM ♂36 | 2.280 | 0.255 |
| | MM ♂25 – MM ♂36 | 1.220 | 1.000 |
| | **SM ♂36 – DM ♂36** | **-6.032** | **<0.001** |
| | **SM ♂36 – MM ♂36** | **-9.182** | **<0.001** |
| | **DM ♂36 – MM ♂36** | **-3.203** | **0.016** |
| | MM ♀25 x ♂25 – MM ♀25 x ♂36 | 2.444 | 0.164 |
| | MM ♀36 x ♂25 – MM ♀36 x ♂36 | -0.094 | 1.000 |
| Contribution of the second and subsequent males to the offspring | **DM ♀25 x ♂25 – DM ♀25 x ♂36** | **-22.693** | **<0.001** |
| | **DM ♀25 x ♂25 – DM ♀36 x ♂25** | **-10.022** | **<0.001** |
| | **DM ♀25 x ♂25 – DM ♀36 x ♂36** | **-12.914** | **<0.001** |
| | DM ♀36 x ♂25 – DM ♀36 x ♂36 | -1.281 | 1.000 |
| | **DM ♀25 x ♂36 – DM ♀36 x ♂36** | **4.382** | **<0.001** |
| | **MM ♀25 x ♂25 – MM ♀25 x ♂36** | **-26.524** | **<0.001** |
| | **MM ♀25 x ♂25 – MM ♀36 x ♂25** | **-6.024** | **<0.001** |
| | **MM ♀25 x ♂25 – MM ♀36 x ♂36** | **-13.264** | **<0.001** |
| | **MM ♀36 x ♂25 – MM ♀36 x ♂36** | **-5.661** | **<0.001** |
| | **MM ♀25 x ♂36 – MM ♀36 x ♂36** | **3.253** | **0.023** |
| | **SM ♀25 x ♂25 – DM ♀25 x ♂25** | **-3.918** | **0.002** |
| | **SM ♀25 x ♂25 – MM ♀25 x ♂25** | **-4.917** | **<0.001** |
| | **DM ♀25 x ♂25 – MM ♀25 x ♂25** | **-5.671** | **<0.001** |

| | | |
|---|---|---|
| **SM ♀25 x ♂36 – DM ♀25 x ♂36** | **-8.268** | **<0.001** |
| **SM ♀25 x ♂36 – MM ♀25 x ♂36** | **-8.806** | **<0.001** |
| **DM ♀25 x ♂36 – MM ♀25 x ♂36** | **-3.261** | **0.022** |
| **SM ♀36 x ♂25 – DM ♀36 x ♂25** | **-4.394** | **<0.001** |
| **SM ♀36 x ♂25 – MM ♀36 x ♂25** | **-4.014** | **0.001** |
| DM ♀36 x ♂25 – MM ♀36 x ♂25 | 1.501 | 1.000 |
| **SM ♀36 x ♂36 – DM ♀36 x ♂36** | **-4.487** | **<0.001** |
| **SM ♀36 x ♂36 – MM ♀36 x ♂36** | **-5.100** | **<0.001** |
| DM ♀36 x ♂36 – MM ♀36 x ♂36 | -2.514 | 0.239 |

**Table S4 – *A posteriori* contrasts of significant explanatory variables for the effect of temperature on mating behavior.** *A posteriori* contrasts with Bonferroni corrections were done to interpret the significant effect of the fixed factors. "T ratio": the T-test value obtained in each comparison. "Latency to the first copulation": time elapsed from pairing until the first mating of a female in seconds; "Duration of the second copulation (remating)": time elapsed from the beginning until the end of the second mating of a female in seconds. Comparisons were made between females developed at 25°C or 36°C ("♀25" and "♀36") and first mated with males developed at 25°C or 36°C ("♂25" and "♂36"). Statistically significant terms ($p \leq 0.05$) are represented in bold.

| Dependent var. | Comparison | T ratio | P-value |
|---|---|---|---|
| Latency to the first copulation | ♀25 x ♂25 – ♀36 x ♂25 | -1.721 | 0.345 |
| | ♀25 x ♂36 – ♀36 x ♂36 | 1.324 | 0.745 |
| | **♀25 x ♂25 – ♀25 x ♂36** | **-7.007** | **<0.001** |
| | **♀36 x ♂25 – ♀36 x ♂36** | **-3.883** | **<0.001** |
| Duration of the second copulation (remating) | **♀25 x ♂25 – ♀36 x ♂25** | **-3.444** | **0.003** |
| | ♀25 x ♂36 – ♀36 x ♂36 | 0.981 | 1.000 |
| | **♀25 x ♂25 – ♀25 x ♂36** | **-5.564** | **<0.001** |
| | ♀36 x ♂25 – ♀36 x ♂36 | -1.177 | 0.966 |

**Supplementary Information 1 – The effect of temperature on the offspring number and sex ratio of females from the mating behavior experiment**

### Statistical analysis

The number of adult offspring and sex ratio were analyzed to infer the degree of female and male fertility, respectively, and to confirm replicability with the results obtained in the previous experiment.

The number of adult offspring was analyzed using a GLMM with a quasi-Poisson error distribution (package *glmmTMB*; Brooks et al., 2017). The sex ratio, calculated as the proportion of daughters over the total number of offspring, was computed using the function *cbind* with the number of daughters and sons as arguments. A GLMM was used with a beta-binomial error distribution and a parameter to account for zero inflation (ziformula ~1; package *glmmTMB*; Brooks et al., 2017). In both models, block, temperature and humidity during mating observations were included as random factors. The developmental temperatures of the female and of the first male (25ºC or 36ºC, in both cases), the female mating history (SM or MM), as well as their interaction, were added as fixed factors.

### Results

The statistical results are summarized in the table below.

| Dependent var. | Explanatory var. | Df | $\chi^2$ | P-value |
|---|---|---|---|---|
| Number of adult offspring | **♀ temperature** | **1** | **169.33** | **<0.001** |
| | ♂ temperature | 1 | 0.229 | 0.632 |
| | ♀ mating history | 1 | 0.983 | 0.322 |
| | ♀ temperature * ♂ temperature | 1 | 0.009 | 0.926 |
| | ♀ temperature * ♀ mating history | 1 | 0.008 | 0.931 |
| | ♂ temperature * ♀ mating history | 1 | 1.513 | 0.219 |
| | ♀ temperature * ♂ temperature * ♀ mating history | 1 | 0.004 | 0.948 |
| Sex ratio | **♀ temperature** | **1** | **22.075** | **<0.001** |
| | ♂ temperature | 1 | 0.370 | 0.543 |
| | ♀ mating history | 1 | 1.023 | 0.312 |
| | ♀ temperature * ♂ temperature | 1 | 1.176 | 0.278 |
| | ♀ temperature * ♀ mating history | 1 | 0.103 | 0.749 |
| | **♂ temperature * ♀ mating history** | **1** | **75.561** | **<0.001** |
| | ♀ temperature * ♂ temperature * ♀ mating history | 1 | 0.079 | 0.779 |

“Df”: the degrees of freedom. “$\chi^2$”: Chi-square value. “Number of adult offspring”: number of offspring that reached adulthood per female; “Sex ratio”: proportion of adult daughters over the total adult offspring. “♀ temperature”: developmental temperature of the females tested; “♂ temperature”: developmental temperature of the first male that mated with a female; “♀ mating history”: the number of times a female mated. Statistically significant terms ($p \leq 0.05$) are represented in bold.

In the figure, panel a) shows the total number of adult offspring thus representing the fertility of the female and panel b) shows the offspring sex ratio (number of daughters/total number of adult offspring), indicating the fertility of the males. Boxplots display a median line, interquartile range (IQR) boxes, 1.5*IQR whiskers and data points. Dark blue, light blue, light orange and dark orange represent treatments where both sexes that were paired were from the control developmental treatment, only the male or the female were heat-stressed, and both sexes were heat-stressed, respectively. White diamonds represent the mean values per condition tested.

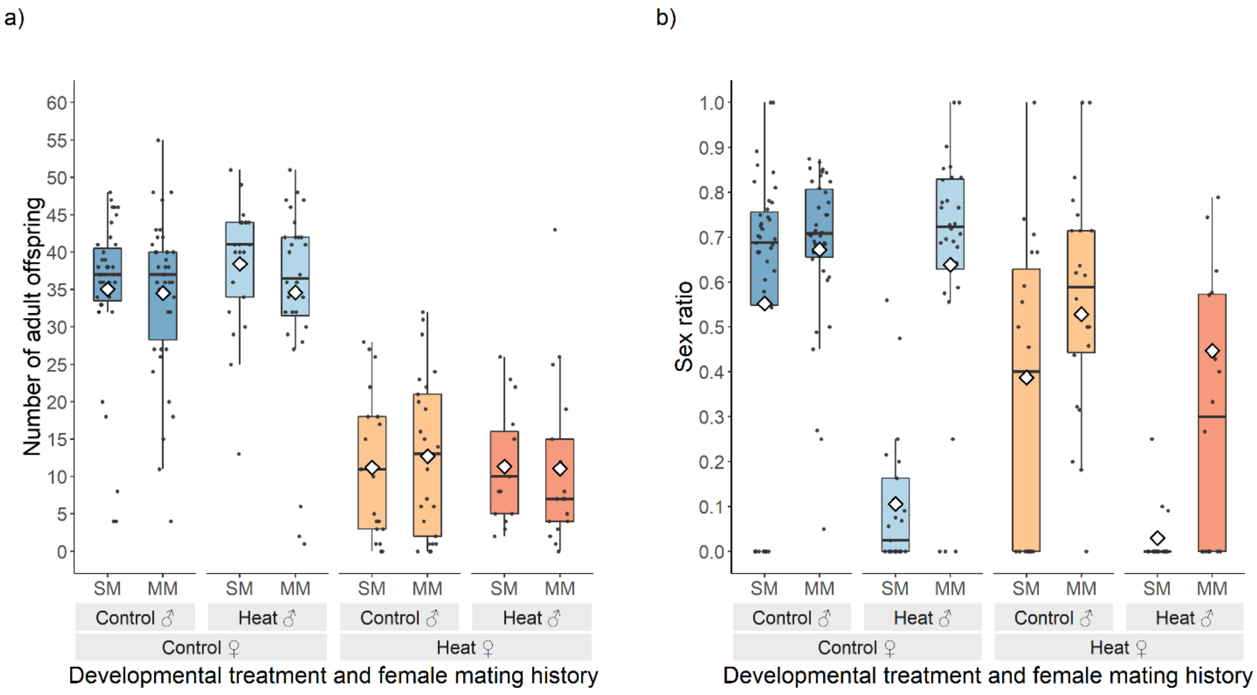


The number of adult offspring was affected by the developmental temperature of the female ($\chi^2_1$ = 169.33, $p$ = <0.001), with heat-stressed females having ca. 67% less offspring compared to control females (12±1.11 offspring compared to 35±0.97), independently of the female mating history and of the developmental temperature of the first male they mated with (panel a).

The offspring sex ratio was affected by the interaction between the developmental temperature of the first male and the female mating history ($\chi^2_1$ = 75.561, $p$ = <0.001). As expected, when first males developed at 25°C, there was no significant effect of the female mating history (SM♂25 *vs* MM♂25: T ratio = 1.011, p = 1.000), with sex ratio averaging 0.60±0.04 (panel b). The proportion of female offspring decreased when the female mated once with a heat-stressed male (0.09±0.05; SM♂25 *vs* SM♂36: T ratio = 9.318, p = <0.001). However, when these females were allowed to remate with control males, the proportion of fertilized offspring increased (SM♂36 *vs* MM♂36: T ratio = -9.076, p = <0.001), reaching values similar to those of females mated with control males (MM♂36 produced ca. 60% daughters; MM♂25 *vs* MM♂36: T ratio = -0.608, p = 1.000). The offspring sex ratio was also affected by the female developmental temperature ($\chi^2_1$ = 22.075, *p* = <0.001), with control females having an overall higher proportion of fertilized offspring than heat-stressed females. These results recapitulate those obtained in the experiment presented in the main text.
